# Designing a hybrid in silico/in-cell controller robust to process–model mismatch associated with dynamically regulated enzyme overexpression

**DOI:** 10.1101/2024.02.27.582404

**Authors:** Tomoki Ohkubo, Yuichi Sakumura, Fuzhong Zhang, Katsuyuki Kunida

## Abstract

Discrepancy between model predictions and actual processes, known as process–model mismatch (PMM), remains a serious challenge in bioprocess optimization. Previously, we proposed the hybrid in silico/in-cell controller (HISICC) concept combining model-based optimization with cell-based feedback to address the PMM problem. Herein, this approach was advanced to regulate intracellular concentrations of rate-limiting enzymes. Advanced HISICC was demonstrated using an engineered *Escherichia coli* strain for fatty acid production (FA3). This strain harbors an in-cell feedback controller, which decelerates acetyl-CoA carboxylase (ACC) overexpression in response to sensing the concentration of malonyl-CoA formed by this enzyme. A mathematical model for FA3 was constructed and validated using experimental data. Simulations assuming various PMM revealed that the HISICC using FA3 effectively mitigates toxicity from excessive ACC by robustly adapting braking its overexpression, minimizing yield loss. This study confirmed HISICC as a viable strategy for enhancing bioprocess efficiency, especially in balancing the bottleneck enzyme levels.

## Introduction

Engineered microorganisms show great promise as cell factories capable of producing a wide array of products, including foods^1^, fuels^2,3^, functional materials^4^, and pharmaceuticals^5^. Achievement of high product yields is crucial for ensuring the economic feasibility of such a technology and is often accomplished *via* the overexpression of bottleneck enzymes of the target production pathways^6^. The tradeoff, however, is that excessive expression of enzymes can adversely affect microbial growth and/or viability^7^. For example, nutrients could be wasted for synthesis of excess enzymes, lowering overall metabolite yields^8–12^. Further, enzyme overexpression may deplete free ribozymes, decreasing cell growth rate. In addition, the accumulation of intermediates and byproducts arising from the enzymatic activity could lead to cytotoxicity^13–17^. Therefore, achieving high-yield production of the desired products necessitates the stringent regulation of enzyme overexpression to maintain appropriate enzyme levels^18–21^. This, in turn, can be achieved *via* two main approaches: model-based process optimization (in silico feedforward controller) and autonomous feedback control by synthetic genetic circuits embedded in microbial cells (in-cell feedback controller)^22–24^.

The in silico feedforward controller functions by adjusting enzyme expression level in response to process inputs such as inducer feeds^8^, temperature^25^, and light^26,27^, which are optimized using mathematical models before the initiation of the bioprocesses. This approach manages the expression of metabolic enzymes *via* the prediction of the future process state, thus maximizing product yields. The challenge, however, arises in the event of a significant process–model mismatch (PMM) between the model prediction and actual process. When PMM occurs, the input values predetermined using the mathematical model would then be suboptimal for the actual process. Model predictive control (MPC) is a method that addresses the constraints arising from PMM, and reoptimizes the process input using measurements from the ongoing process in real time^28–31^. Although MPC as a control strategy offers robustness and optimal control in the event of PMMs, monitoring the cellular state, including intracellular concentrations of RNA, metabolites, and enzymes, presents its own set of challenges.

The in-cell feedback controller, emerged from synthetic biology, detects the concentrations of intracellular metabolites^32–34^, extracellular nutrients^35^, or cell density^9,36,37^, and regulates enzyme expression according to these signals. While the in silico feedforward controller is effective in maximizing product yield, the in-cell feedback controller can detect intracellular states that are difficult to monitor with process sensors or biochemical analyses and provides feedback on enzyme expression *in situ*. Therefore, the in silico feedforward and in-cell feedback controllers complement each other with respect to their limitations.

We previously proposed a hybrid control system (hybrid in silico/in-cell controller, HISICC) that combined an in silico feedforward controller with an in-cell feedback controller to overcome the PMM-associated difficulties inherent to model-based process optimization^38^. We designed and evaluated a HISICC for resource allocation as a proof-of-concept; the HISICC contained an in-cell feedback controller that autonomously triggered the overexpression of enzymes participating in an alcohol biosynthesis pathway concomitantly with the shutdown of a competing pathway for cell growth based on quorum sensing. The concept of HISICC showed great promise as a solution to PMM-associated difficulties; however, the in-cell feedback control of cell density that relied on quorum sensing is amenable to substitution with electronic control using a spectrophotometer. Hence, an important advantage of our approach—the dynamic adaptation of enzyme overexpression based on the *in situ* sensing of intracellular metabolites or intermediates, a challenging prospect using other means such as biochemical sensors or analysis techniques—remains to be demonstrated.

In the current study, we sought to demonstrate the advantage of our approach for achieving regulated overexpression of rate-limiting enzymes using the engineered *Escherichia coli* strains FA2 and FA3, originally developed by Liu *et al*. for fatty acid production^39^, as well as the strain FA4, conceptually designed based on the strains FA2 and FA3. The overexpression of the bottleneck enzyme acetyl-CoA carboxylase (ACC) in these strains is amenable to external tuning using isopropyl β-D-thiogalactopyranoside (IPTG). As detailed in the Results section, prediction errors in intracellular ACC concentrations constitute a critical PMM in this process and lead to lower fatty acid yields. The FA3 strain contains an in-cell feedback controller that indirectly detects increase in ACC concentrations; this strain therefore forms a HISICC when combined with an in silico feedforward controller (Figure 1), whereas the other two strains form no-feedback systems. First, the three strains were modeled to allow the design of corresponding in silico controllers. Next, the robustness of the HISICC in the event of PMM was evaluated by comparing the fatty acid yields obtained with the HISICC versus the no-feedback counterparts in multiple rounds of simulation assuming PMMs of various magnitudes. The results indicated that our approach could mitigate the yield loss attributable to PMMs associated with the regulation of enzyme overexpression.

**Figure 1.**
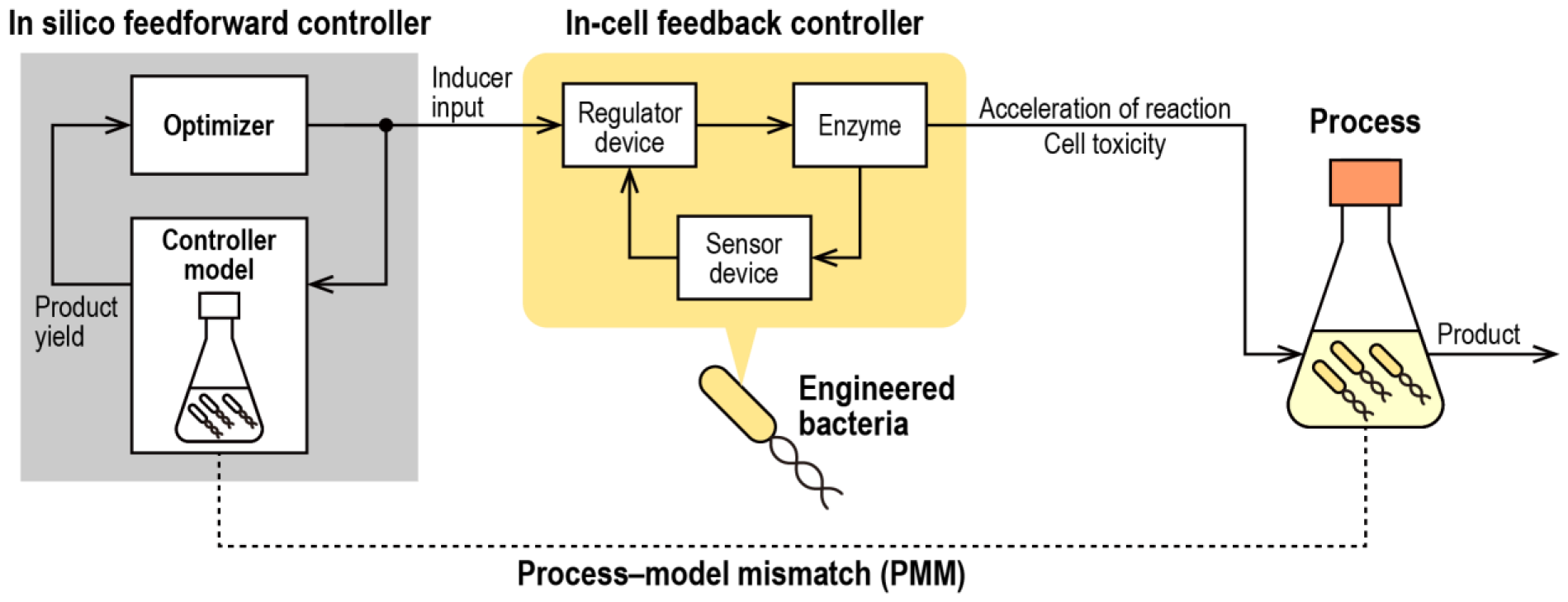
Conceptual diagram of the hybrid in silico/in-cell controller (HISICC). The in silico feedforward controller optimizes the inducer input to maximize the product yield using the controller model by balancing the acceleration of bottleneck reactions with the cytotoxicity caused by overexpression of the rate-limiting enzyme. The in-cell feedback controller in the bacterial cells autonomously performs feedback control of the enzyme concentration in response to the optimized inducer input.

## Results

### Fatty acid production in engineered *E. coli* strains

With a view to facilitate a better understanding of the mathematical modeling, we have provided herein an overview of the two engineered *E. coli* strains FA2 and FA3 that were previously developed by Liu *et al*.^39^, as well as the strain FA4 that was conceptually designed for simulation in the current study. The first step in the fatty acid biosynthesis pathway in *E. coli* involves the conversion of acetyl-CoA to malonyl-CoA by ACC, encoded by *accABCD*. This reaction is the rate-limiting step of the pathway^15^, and several studies have shown that ACC overexpression increases both intracellular malonyl-CoA concentration and fatty acid yields^13,40^. However, the overexpression of ACC is toxic to cells; the exact molecular mechanism underlying the toxicity is currently unclear, but excessive ACC has been reported to decrease cell growth^13–15^ as well as fatty acid production per unit cell density^39^. Therefore, an ACC concentration that is either too high or too low reduces the fatty acid yield of the bioprocess, which reflects the tradeoff associated with ACC overexpression. The maximization of fatty acid yield requires a rapid stabilization of the ACC concentration to an optimal level, offering a balance between cytotoxicity and improvements in the reaction rate.

The *E. coli* strain FA2 carries the regulator device, a genetic device containing a plasmid copy of *accABCD* under the control of the T7 RNA polymerase, allowing the ACC overexpression to be externally controlled using IPTG (Figure 2A and 2B). Thus, in the bioprocess using this strain, IPTG concentration is defined as the input variable to be optimized by the in silico feedforward controller. Unlike the other two strains, this strain lacks mechanisms for the reduction of ACC overexpression during midculture period. Therefore, to alleviate the toxicity of ACC, the in silico feedforward controller coupled to this strain needs to maintain the initial ACC expression at a low level to ensure that ACC concentrations in the latter half of the culture period do not become too high (Figure 2C).

**Figure 2.**
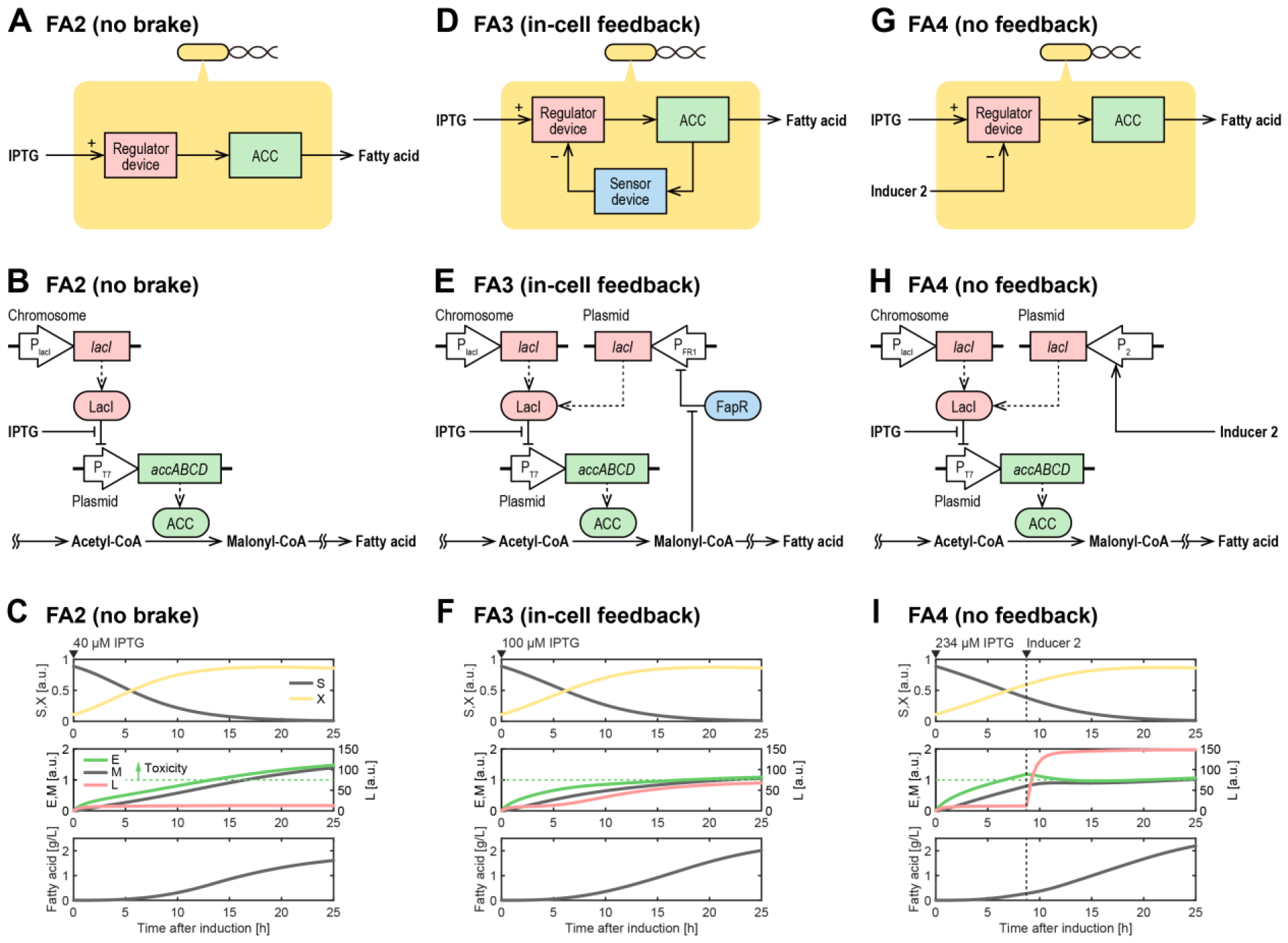
Designs and functions of the engineered *E. coli* strains. (Top row) Diagrammatic representation of the control circuits in the *E. coli* strains FA2 (A), FA3 (D), and FA4 (G). The regulator device is designed to allow ACC overexpression to be externally tunable using IPTG, and the sensor device, to detect increase in ACC concentration by sensing malonyl-CoA levels. The arrows with plus (+) or minus (−) signs represent upregulation or downregulation, respectively. (Middle row) Synthetic genetic circuits in the *E. coli* strains FA2 (B), FA3 (E), and FA4 (H). The boxes with rounded edges and dotted arrows represent proteins and protein expression, respectively. The red and blue boxes correspond to the regulator and sensor devices, respectively. (Bottom row) Simulated dynamics of the state variables in the *E. coli* strains FA2 (C), FA3 (F), and FA4 (I). The symbols for the variables are listed in Table 1. The horizontal green dotted line represents the toxicity threshold of ACC *E*_tox_; when ACC concentration *E* exceeds *E*_tox_, the ensuing toxicity compromises fatty acid secretion per unit cell density.

The *E. coli* strain FA3 contains a genetically encoded feedback controller consisting of the regulator device and another genetic device, the sensor device (Figure 2D). The sensor device is composed of FapR, a malonyl-CoA-responsive transcription factor, and FR1, a FapR-regulated promoter (Figure 2E). In this design, if ACC level is too high, the produced malonyl-CoA will trigger the sensor device to express LacI, which in turn represses ACC overexpression. Meanwhile, the in-cell feedback controller can be externally initiated by IPTG, which alleviate the repression of ACC overexpression by LacI. Overall, the in-cell feedback controller integrated in the FA3 strain indirectly detects the increase in ACC concentration by sensing malonyl-CoA levels and responds by reducing ACC expression during the midculture period. Therefore, the in silico feedforward controller coupled to this strain can employ higher IPTG concentrations for inducing the cells, leading to higher initial expression of ACC as compared to the in silico controller coupled to the strain FA2; this, in turn, shortened the response time for achieving the optimal ACC concentration (Figure 2F). In fact, 27% higher yield of fatty acids was obtained with the strain FA3 as compared to the strain FA2 under identical culture conditions (with the exception of IPTG concentration employed for induction) in the previous study^39^.

**Table 1.** State and input variables for the models.

| Symbol | Initial value | Unit | Description |
| --- | --- | --- | --- |
| $S$ | $1 - X(0)$ | Dimensionless | Extracellular substrate concentration |
| $X$ | Estimated | Dimensionless | Normalized cell density |
| $E$ | 0 | Dimensionless | Intracellular ACC concentration |
| $M$ | 0 | Dimensionless | Intracellular malonyl-CoA concentration |
| $L$ | Estimated | Dimensionless | Intracellular LacI concentration |
| $P$ | 0 | Dimensionless | Extracellular fatty acid concentration |
| $I$ | — | $\mu\text{M}$ | Extracellular IPTG concentration (input) |
| $u$ | 0 | Dimensionless | Inducer 2 input for *E. coli* strain FA4, $u$ is either 0 or 1 (unit step function) |

The *E. coli* strain FA4 was conceptually designed based on the strains FA2 and FA3. This strain exhibits reduction in ACC overexpression in response to a second inducer (Inducer 2) added during the midculture period. The Inducer 2 induces the expression of an additional *lacI* downstream Promoter 2, an Inducer 2–regulated promoter (Figure 2G, 2H, and 2I). Either the arabinose/*araBAD* promoter or rhamnose/*rhaBAD* promoter pairs^41^ could be employed as the inducer/promoter pair for proof-of-concept and the actual development of this strain, with both IPTG and Inducer 2 defined as process inputs for the bioprocesses employing this strain. Among the three strains employed herein, only the strain FA3 contains an in-cell feedback controller; therefore, only the coupling of the strain FA3 with the in silico feedforward controller is regarded as a HISICC.

### Mathematical modeling

#### FA2 strain model

This model forms the basis for modeling the other two strains; the other two models are extensions of this one, with the addition of a term representing LacI expressed from plasmids to a differential equation for LacI dynamics. This model consists of two differential equations describing substrate consumption and cell growth and four differential equations describing the normalized concentrations of important compounds in the cell.

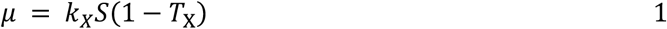

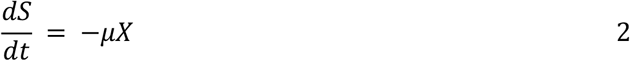

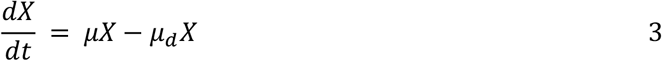

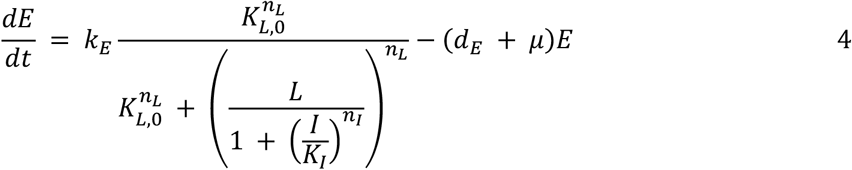

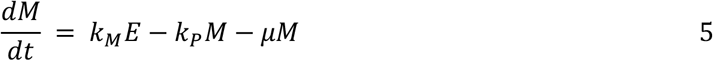

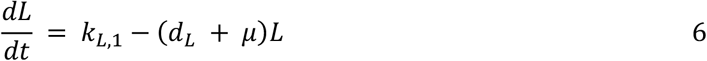

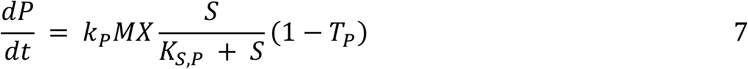

The descriptions and initial values of the six state variables are summarized in Table 1, and the model parameters are listed in Table 2.

**Table 2.**
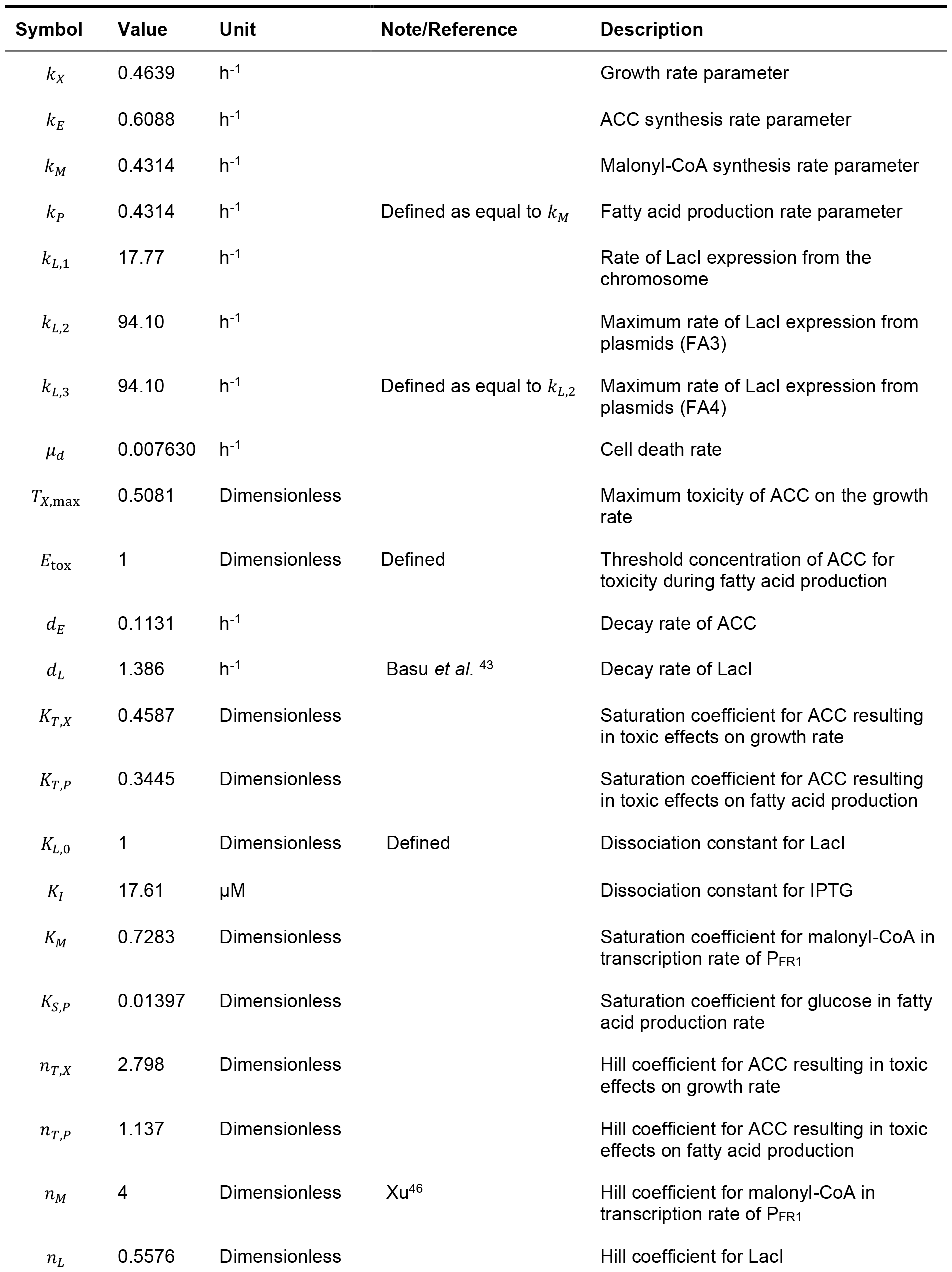
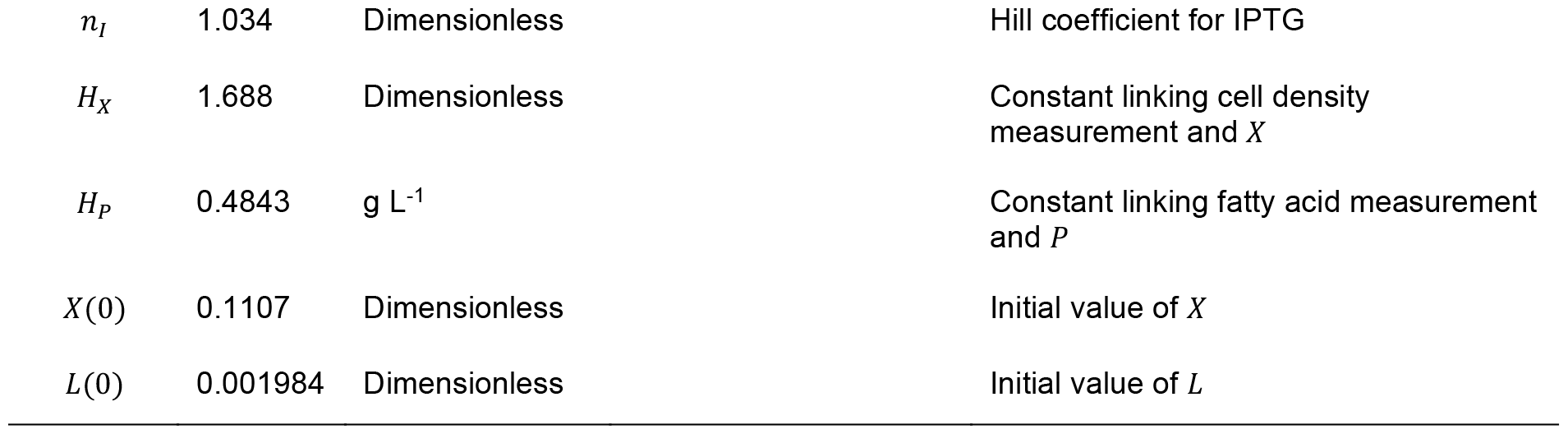
Model parameters. The parameter values have been estimated for all datasets, except those defined or adapted from literature.

The equations for cell growth were derived based on Monod’s model^42^, with some modifications (Equations 1–3). First, the saturation of the specific growth rate μ at high substrate concentrations was ignored to simplify the model. Second, cell death was assumed to occur at a constant rate μ_*d*_. *T*_X_ represents the deceleration in cell growth owing to ACC cytotoxicity, as described below.

Equations 4–6 describe the dynamics of the intracellular concentrations of ACC (*E*), malonyl-CoA (*M*), and LacI (*L*), respectively. These three compounds were assumed to be diluted upon cell growth. The concentrations of ACC and LacI were also assumed to be reduced because of protein degradation. The LacI degradation rate *d*_*L*_ was adopted from Basu *et al*.^43^ The expression of ACC is regulated by the T7 promoter and is amenable to tuning using various concentrations of IPTG. The mathematical expression for the promoter output as a function of both LacI and IPTG concentrations was adopted from Gardner *et al*.^44^ (Equation 4). The saturation constant *K*_*L*,0_ was defined as 1, indicating that *L* is normalized with respect to *K*_*L*,0_. The production and consumption rates of malonyl-CoA were assumed to be proportional to the concentrations of ACC and malonyl-CoA, respectively (Equation 5). By defining the proportionality constants *k*_*M*_ and *k*_*P*_ as equal, we equalized the ranges of the nondimensionalized state variables *M* and *E*. LacI was assumed to be expressed at a constant rate *k*_*L*,1_ from the chromosomal *lacI* gene (Equation 6).

Equation 7 describes the fatty acid concentration in the culture medium. The volumetric fatty acid production rate was assumed to be proportional to both the cell density *X* and the specific malonyl-CoA consumption rate *k*_*P*_*M*. The saturation constant *K*_*S,P*_ was introduced to reproduce the slowdown of fatty acid production per unit cell density due to the depletion of substrates in the culture medium. *T*_*P*_ represents the slowdown of fatty acid production due to the cytotoxicity associated with ACC overexpression, as discussed in detail below.

This model assumes that excessive ACC concentration affects the specific rates of cell growth as well as fatty acid production.

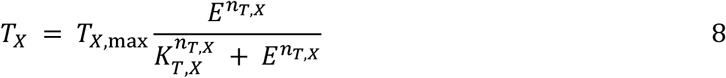

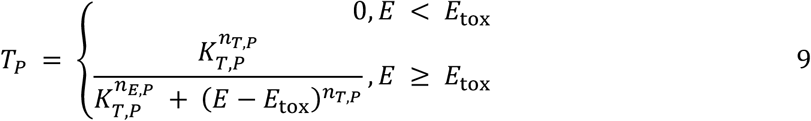

*T*_*X*_ represents the toxic effects on cell growth; an increase in *E* leads to increase in *T*_*X*_ and decrease in the specific cell growth rate μ (Equations 1 and 8). *T*_*P*_ represents the toxic effects on fatty acid production. We have assumed that when *E* exceeds the threshold *E*_tox_, *T*_*P*_ increases and fatty acid production decelerates (Equations 7 and 9). The value of *E*_tox_ was defined as 1 herein, indicating that *E* is normalized with respect to *E*_tox_.

Equation 10 is an observation equation that links the model simulations to the experimental measurements. *y*_1_ and *y*_2_ represent the relative cell density, calculated on the basis of optical density (OD_600_) and fatty acid concentration, respectively. *H*_*X*_ and *H*_*P*_ are proportionality constants.

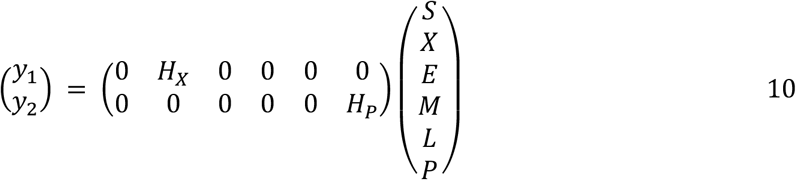

#### FA3 strain model

The strain FA3 contains an in-cell feedback controller that senses an increase in malonyl-CoA concentration and promotes the overexpression of LacI from plasmids to reduce ACC expression as needed. A mathematical representation of this feedback was generated by extending the FA2 strain model; we added a term representing *lacI* expression from plasmids to the differential equation describing LacI dynamics in Equation 6.

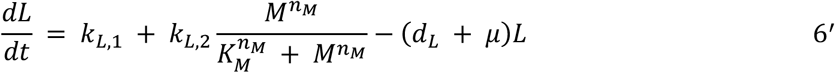

The response of the FR1 promoter to malonyl-CoA was approximated using Hill’s equation. The Hill coefficient *n*_*M*_ was defined as 4, in accordance with the modeling of another in-cell feedback controller for fatty acid production that employed the same FapR-based malonyl-CoA sensor as the strain FA3^45,46^. To simplify the model, the concentration of FapR was assumed to be constant and excluded from the model. The other equations and parameter values were shared with the FA2 strain model.

#### FA4 strain model

The strain FA4 contains a brake circuit designed to decelerate ACC overexpression by promoting the overexpression of LacI from plasmids in response to Inducer 2. A mathematical representation of this brake function was generated by adding a term representing *lacI* expression from plasmids to Equation 6 in the FA2 strain model.

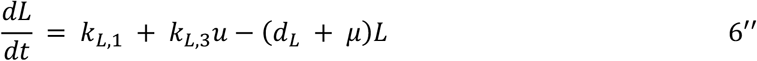

*u* represents the normalized concentration of Inducer 2. We have defined *u* = 1 as the concentration of Inducer 2 at which a saturation of Promoter 2 response is attained. The maximum *lacI* expression rate *k*_*L*,3_ was arbitrarily defined as equal to *k*_*L*,2_ in the FA3 strain model. The other equations and parameter values were shared with the FA2 strain model.

### Model simulation and validation

The FA2 and FA3 strain models were trained using experimental data from fatty acid production cultures conducted in a previous study^39^. Details pertaining to the experimental data have been presented in the Methods section. As aforementioned in the mathematical modeling subsection, the two models share most equations and parameters; therefore, we trained the two models simultaneously using experimental data from the two strains collectively as a single training dataset. In other words, a single value was estimated for each parameter common to the two models such that the two models could simultaneously fit the corresponding experimental data. Trained using all available experimental data, both the models exhibited close fit with the experimental data (Figure 3 and Table 2). This, in turn, suggests that despite their simple structure, the models successfully captured the dynamics of cell growth and fatty acid production by the two strains in response to various IPTG inputs. Further, the holdout validation method (described in detail in the Methods section) was employed to ensure that the trained models did not overfit the training dataset (Figure 4). FitPercent, the normalized root-mean-squared error expressed as a percentage^47^, was employed as an indicator to determine how well the model response fit the validation dataset. FitPercent of >70% was obtained for all the validation datasets, indicating that the model did not overfit the training data and had good generalization performance within the range of IPTG concentrations applied in the experiments.

**Figure 3.**
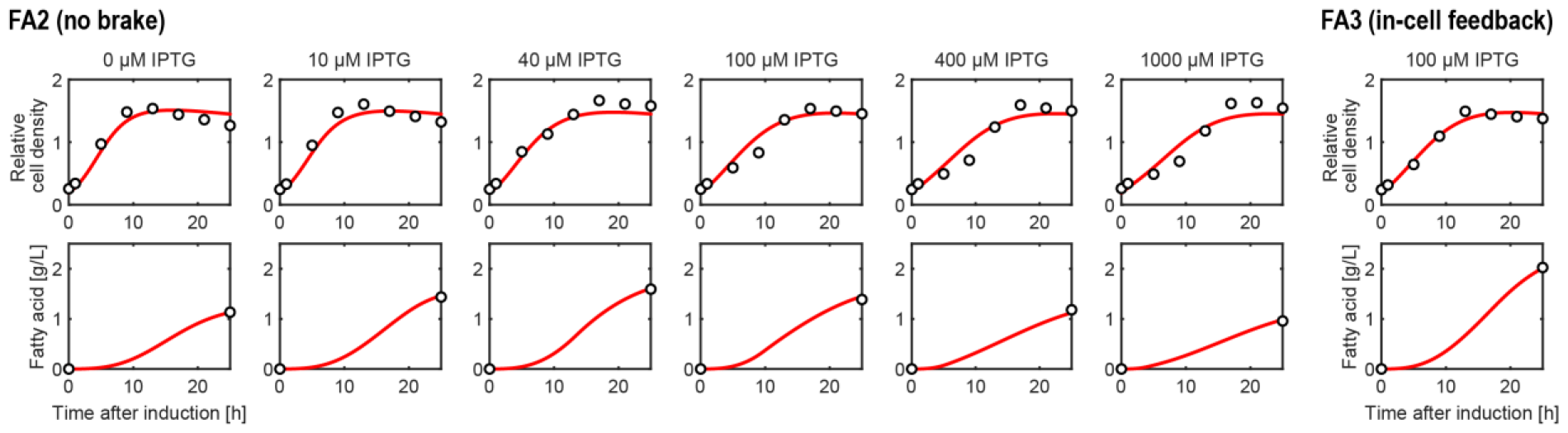
Simulation of fatty acid production by the *E. coli* strains FA2 or FA3 upon induction with various concentrations of IPTG. The models were trained using all available datasets. The white dots and red lines represent the experimental data and simulations, respectively.

**Figure 4.**
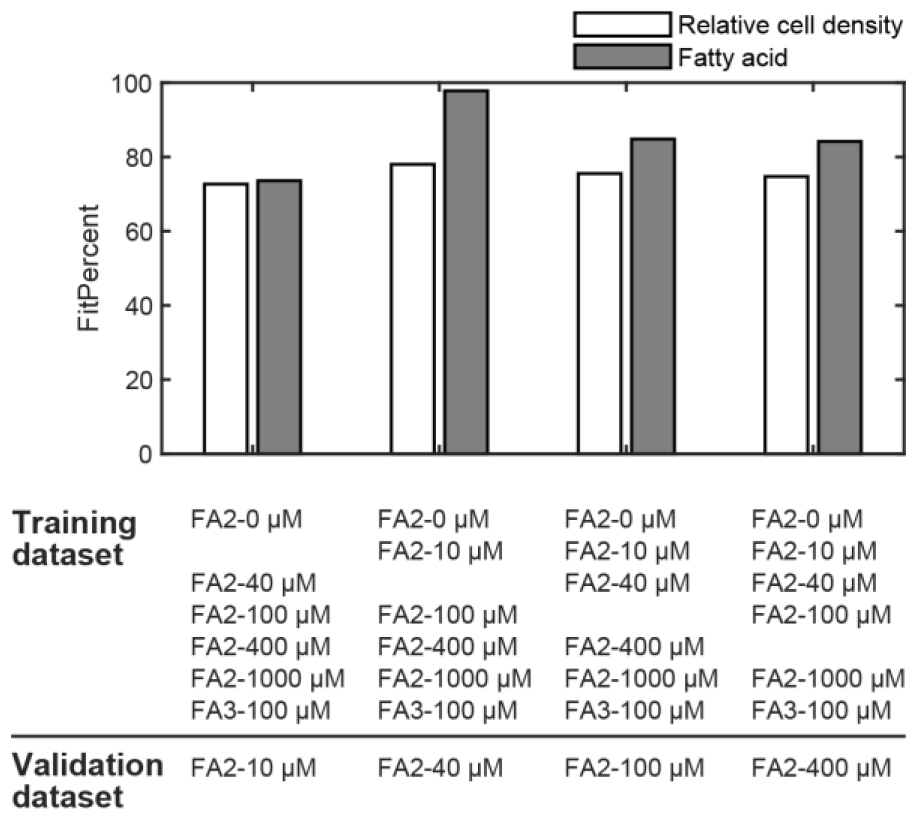
Holdout validation of the FA2 and FA3 strain models. Validation was performed separately for cell density and fatty acid concentration. The normalized root-mean-squared error expressed as a percentage (FitPercent) was used as an indicator of model validation. In each round of validation, data obtained from one batch of the FA2 strain were employed as the validation dataset, and data from the other batches of FA2 and FA3 strains, as the training dataset.

### Model-based input optimization

We subsequently sought to demonstrate the performance of the in silico feedforward controllers in the HISICC (strain FA3) or no-feedback (strains FA2 or FA4) systems; toward this end, we optimized the IPTG input to obtain maximum fatty acid yields. The FA2 and FA3 strain models were trained using all experimental datasets prior to input optimization (Figure 3, Table 2). The FA4 strain model shared all parameters with the other two models and had no unique parameters, negating any necessity for training it. The in silico feedforward controllers for the strains FA2 and FA3 optimized the IPTG concentrations (Figure 5A and 5B), while the controller for the strain FA4 simultaneously optimized both IPTG concentration and the time of addition of Inducer 2 (Figure 5C). The feasible input range, as per the experimental conditions, was defined as 0–1000 μM for IPTG concentration and 0–25 h for the time of addition of Inducer 2. Furthermore, a comprehensive simulation of the models was carried out to visualize the overall distribution of fatty acid yields over the feasible input range. The predicted fatty acid yields for all the strains exhibited a convex-upward function with a distinct peak, and the input values optimized by the in silico controllers were consistent with this peak. This observation suggests that the in silico feedforward controllers contribute to product yield improvement through input optimization. In addition, the optimal IPTG concentration for the strain FA2 was the lowest among the three strains; this observation is consistent with the inability of the strain FA2 to reduce ACC expression during the midculture period unlike the other two strains, requiring the in silico controller to maintain a lower initial IPTG concentration to maintain ACC expression at a low level initially. This, in turn, suggests that the models successfully reproduce the tradeoff inherently associated with ACC overexpression for fatty acid production.

**Figure 5.**
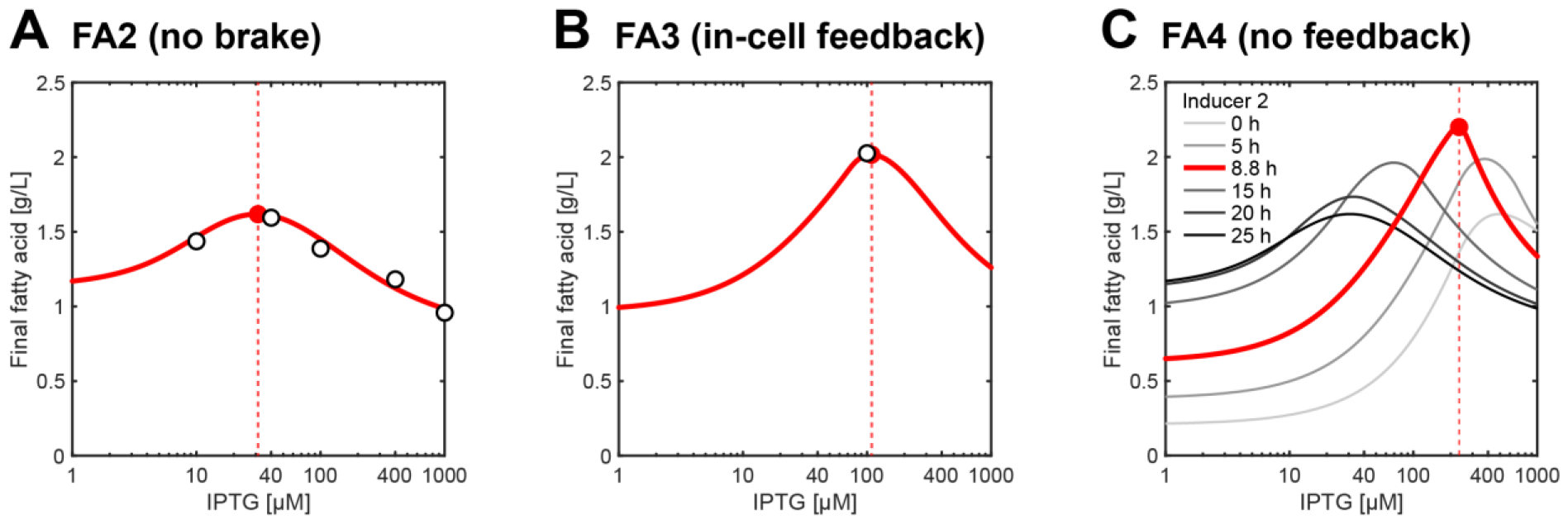
Model-based optimization of the IPTG input to achieve maximum fatty acid yields. The white dots and red lines represent the experimental data and simulations, respectively. The red dots and dotted vertical lines represent the optimized IPTG concentrations for the *E. coli* strains FA2 (A), FA3 (B), and FA4 (C). The gray lines represent simulations with Inducer 2 added at various times. The red line (C) represents the simulation with the optimal time of addition of Inducer 2.

### Controller performance against PMM

The robustness of the HISICC against PMM was subsequently evaluated by calculating the fatty acid yields for the three strains using a multiround simulation (Figure 6). For all the simulation rounds, the parameters in the controller models for the strains FA2 and FA3 were estimated using all experimental datasets (Figure 3, Table 2); these parameter estimates were also employed in the controller model for the strain FA4. Thus, the inducer input values employed in this simulation were identical to the optimized values (described in the Model-based input optimization subsection; Figure 5, dotted vertical lines). Each simulation round introduced PMMs of different magnitudes, and ACC expression in the strains was assumed to be faster or slower than the values predicted by the corresponding in silico feedforward controllers. In other words, we computationally reproduced the common scenario in which enzyme overexpression deviates from the values predicted by the controller. Specifically, we defined various values for the parameter *k*_*E*_ in Equation 4, which represents the maximum rate of ACC expression, for each round to calculate the dynamics of the process; these values were denoted as 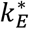 to distinguish them from the corresponding value *k*_*E*_ in the controller model. Notably, 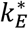 represents an intrinsic property of the cells that is difficult to manipulate in cell culture–based experiments. The round with 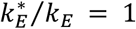 corresponds to the situation in which no PMM was introduced (Figure 6; dotted vertical line). In other words, this round represents the ideal situation in which the in silico feedforward controllers accurately predicted the ACC dynamics.

**Figure 6.**
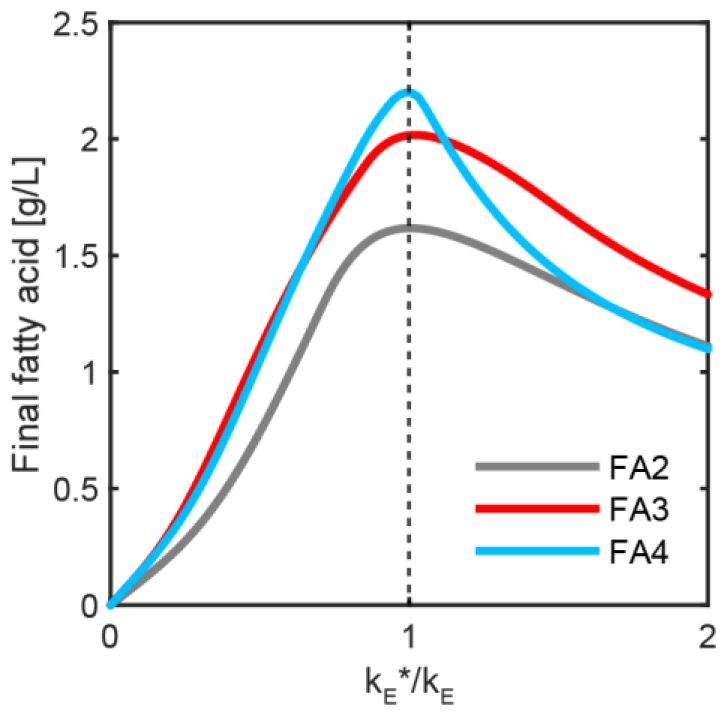
Fatty acid yields under conditions of PMMs of various magnitudes were calculated in a multiround simulation. 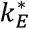 represents the maximum rate of ACC expression set to calculate the process dynamics in each simulation round. *k*_*E*_ represents the value set for the same parameter in the controller model, which was set to the estimated value listed in Table 2 for all rounds of simulation. Consequently, the inducer inputs (IPTG concentration for the strains FA2 and FA3; IPTG concentration and time of addition of Inducer 2 for the strain FA4) were also set for all rounds at the optimized values shown in Figure 5. The dotted vertical line represents a simulation round with no PMM (i.e., the same value was set for the parameter in the controller model and in the model used to simulate the actual process).

Highest fatty acid yields were obtained in the no-PMM round with all three strains. Lower yields were obtained when ACC expression was slower than predicted 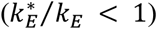, apparently because sufficient ACC accumulation was not attained during the 25-h culture period. Similarly, a faster-than-predicted expression of ACC 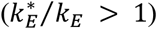 also resulted in lower yields of fatty acids because of the cytotoxicity associated with ACC overexpression.

Among the three strains, the lowest yield of fatty acids was obtained with the strain FA2, regardless of 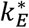. This is attributable to the inability of the strain FA2 to decelerate ACC expression during the midculture period, unlike the other two strains. The consequent accumulation of excessive ACC impaired fatty acid production in the later stages of the culture.

Differences were observed between the strains FA3 and FA4 with respect to interference by PMMs even though both the strains were designed to allow braking of ACC overexpression. Under conditions of slow rates of ACC expression 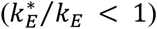, little difference was observed in the fatty acid yields between the two strains. Under the conditions when ACC expression was comparable to that predicted by the controller 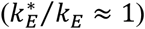, the strain FA4 produced slightly higher quantities of fatty acids than the strain FA3; this is because the in silico controller for the strain FA4 could induce ACC overexpression with a higher concentration of IPTG and subsequently resort to abrupt braking enabled by the input of Inducer 2 (Figure 2I). Such abrupt braking is impossible for the strain FA3 because the in-cell feedback controller embedded in this strain exhibits a gradual response to increase in malonyl-CoA concentrations (Figure 2F). Interestingly, under conditions of rapid rate of ACC expression 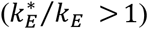, the strain FA4 produced lower yields of fatty acids than the strain FA3. The reasons underlying this difference in robustness to the PMM are elucidated below. The in silico feedforward controller for the strain FA4 predetermined the time of addition of Inducer 2, independent of the actual ACC concentration; a faster accumulation of ACC than that predicted by the controller was obtained. Consequently, the time of addition of Inducer 2 was later than the actual optimal timing, allowing excess ACC accumulation and thereby, reduced fatty acid yields. In contrast, the strain FA3 contains the in-cell feedback controller, allowing an autonomous change in the timing and intensity of brake application on ACC overexpression in accordance with the observed increase in malonyl-CoA concentration (reflecting ACC activity). Thus, a reduced loss of fatty acid yield was obtained for the strain FA3 than for the strain FA4. Additionally, under conditions of very fast rate of ACC expression 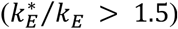, the braking effect triggered by Inducer 2 was almost nonexistent, resulting in low fatty acid yields nearly equivalent to that of the strain FA2 that lacks any brake circuit. These results indicate that within the HISICC, the in-cell feedback controller can support the in silico feedforward controller to mitigate issues arising from the PMM associated with the overexpression of the key enzyme.

## Discussion

The current study elucidates the extension of the HISICC concept, which we previously devised to resolve the PMM issue^38^, to a system for regulating the overexpression of a rate-limiting enzyme. The engineered *E. coli* strains FA2 and FA3, developed by Liu *et al*. for fatty acid production^39^, were employed for a proof-of-concept study. The strain FA3 has an in-cell feedback controller that indirectly detects an increase in the concentration of the rate-limiting enzyme by sensing an intermediate produced by the enzymatic activity; it then applies a brake on the expression of the enzyme, unlike the strain FA2. Therefore, only the strain FA3 (and not FA2) forms a HISICC in combination with an in silico feedforward controller. We hypothesized that the in-cell feedback controller allows the HISICC to adjust the timing and strength of the brake application as per the detected enzyme concentration; this, in turn, allows a reduction in fatty acid yield loss attributable to the PMM associated with the rate of enzyme expression. As a first step to prove this hypothesis, we constructed mathematical models and designed in silico feedforward controllers for the strains. Trained using experimental data, the models captured the dynamics of cell growth and fatty acid production in response to various inducer inputs. They also exhibited good predictive performance against the experimental data during holdout validation. Next, we evaluated the robustness of the HISICC against the PMMs using a multiround simulation. We compared the strains FA2, FA3, and FA4; the strain FA4 was conceptualized based on the circuit designs in the strains FA2 and FA3, and contains a brake circuit that decelerates the rate of enzyme overexpression in response to another inducer. A comparison of fatty acid yields from the three strains revealed that when the rate of enzyme overexpression was higher than predicted by the in silico controllers, the minimum fatty acid yield loss was obtained with the strain FA3 containing the in-cell feedback controller, which is in line with our hypothesis. This observation demonstrates the efficacy of the HISICC as a unique solution for PMM-associated issues during the overexpression of rate-limiting enzymes.

The experimental data from only one batch of the strain FA3 were available for parameter estimation; the lack of experimental data for the strain FA3 was circumvented by taking advantage of the fact that most of the genetic circuits were common to the strains FA2 and FA3. The two strains were modeled such that most of the equations and parameters were shared, and a single training dataset was prepared by combining the data from both the strains. As a result, despite having only two unique estimated parameters (*k*_*L*,2_ and *K*_*M*_), the trained model of the strain FA3 fitted the experimental data and consequently reproduced the improvement in fatty acid yields compared to the strain FA2 (Figure 3). This suggests that both the FA2 and FA3 strain models have sufficient prediction performance as the controller models for the corresponding in silico controllers. The small size of the training dataset resulted in long confidence intervals relative to the estimated values for some parameters (data not shown); however, in general, the prediction performance of the trained model is independent of the length of the confidence interval of the model parameters, and thus, so is the demonstration of the HISICC in this study.

During modeling of the strain FA4, we arbitrarily defined the maximum strength *k*_*L*,3_ of Promoter 2 to be equal to *k*_*L*,2_ of the FR1 promoter used in the strain FA3. Although the strain FA4 is only conceptual as of now, the construction of this strain is possible using various inducible promoters as Promoter 2, as noted in the subsection on engineered *E. coli* strains (Figure 2H). Depending on the choice of the promoter, the value of *k*_*L*,3_ could be smaller, leading to a weaker braking effect on ACC overexpression and lower fatty acid yields as compared to the simulation presented in this study. However, this does not affect our argument that the HISICC using the strain FA3 exhibits greater robustness against the PMMs associated with the rate of enzyme expression than the no-feedback system using the strain FA4.

In the current study, the in silico feedforward controllers were designed to optimize only one or two inputs: the IPTG concentration for the strains FA2 and FA3, and IPTG concentration as well as time of addition of Inducer 2 for the strain FA4. These input values could also be optimized by exhaustive experiments. However, if multiple rate-limiting enzymes need to be regulated, as with another *E. coli* strain engineered for fatty acid production by Xu *et al*.^45^, the cost of input optimization by exhaustive experiments is likely to be prohibitively high. Moreover, the induction of genes using light^11,26,27^, temperature^25^, or consumable inducers such as glucose^48^, lactose^49^, and arabinose^50^ is reversible and allows for time-varying inputs to the in-cell feedback controller. Although these reversible input channels enable more effective and dynamic regulation of the expression of key enzymes, they require determination of the optimal input values for each control interval; conducting exhaustive experiments would therefore be very challenging. Thus, the HISICC concept demonstrated in this study is expected to lead to a practical solution to the PMM-associated issues in more complicated bioprocesses involving the manipulation of multiple enzymes or time-varying process inputs.

## Methods

### Experimental data

The experimental data from fatty acid production cultures used to train and validate the FA2 and FA3 strain models were obtained from Liu *et al*.^39^ (Figure S2 and S4 in the Supplementary Information appended to the original article). We provide herein, a brief description of the fatty acid production culture conditions. For both strains, the seed cultures were grown overnight at 37°C on a rotary shaker at 220 rpm in Luria–Bertani medium supplemented with the appropriate antibiotics (50 mg/L ampicillin, 50 mg/L kanamycin, and 30 mg/L chloramphenicol). The seed cultures were then transferred, using 2% (v/v) inoculation, to minimal medium (M9 minimal medium supplemented with 75 mM MOPS, 2 mM MgSO_4_, 1 mg/L thiamine, 10 μM FeSO_4_, 0.1 mM CaCl_2_, micronutrients including 3 μM [NH_4_]_6_Mo_7_O_24_, 0.4 mM boric acid, 30 μM CoCl_2_, 15 μM CuSO_4_, 80 μM MnCl_2_, and 10 μM ZnSO_4_) supplemented with 2% glucose and the appropriate antibiotics, and grown overnight for adaptation. Fatty acid production cultures were initiated from the overnight culture for adaptation with an initial OD_600_ of 0.08 and grown in fresh minimal medium supplemented with 2% glucose and 0.01% arabinose. When an OD_600_ of 0.6 was attained, the strains were induced with 200 nM anhydrotetracycline and various amounts of IPTG. Relative cell density (arbitrary units) was defined in the original work^39^ using the OD_600_ values recorded every 1000 s. We decimated the time-series data of relative cell density into 4-hourly datasets for parameter estimation. The samples for fatty acid quantification were collected 25 h after induction.

### Parameter estimation

MATLAB/Simulink 2022a was employed for model construction and simulation. Simulink Design Optimization was used for estimating the model parameters. The parameter values were chosen to minimize the sum of the squared errors between the model predictions and measured data, as shown in Equations 10 and 11. Errors were normalized to the maximum values of measurements in the same culture. In Equations 10 and 11, *V* represents the objective function for optimization. The vectors ***θ*** and 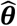 represent the model parameters and their estimated values, respectively. *y* and *ŷ* represent the measured and predicted process outputs, respectively. *I* represents the IPTG input. The subscripts *i, j, k*, and *l* represent the process output (*i* = 1 for relative cell density and *i* = 2 for fatty acid concentration), strain (*j* = 1 for FA2 and *j* = 2 for FA3), IPTG input, and measurement time indices, respectively. *N*_*i*_ and *M*_*j*_ represent the number of measurement time points for the process output *i* and the number of IPTG input levels for the strain *j*, respectively. The composition of the datasets and the corresponding indices are summarized in Tables S1 and S2.

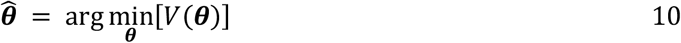

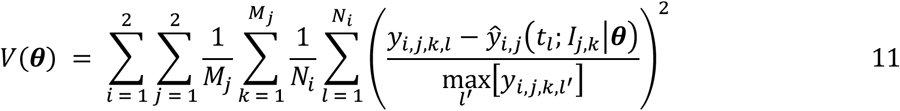

The *lsqnonlin* command was used for optimization. The trust region method was selected as the optimization algorithm for this command. A scaling factor was specified for each parameter to prevent large absolute values from excessively influencing the overall parameter estimation.

### Model validation

The holdout method was employed for validation that the models predicted experimental results from various IPTG concentrations without overlearning. For each round of validation, one batch of the strain FA2 was selected from six batches, excluding the batches induced at the highest (1000 μM) or lowest (0 μM) IPTG concentrations; the data obtained from this batch were defined as the validation dataset (Figure 4). Data obtained from the remaining five batches of the strain FA2 and one batch of the strain FA3 were collectively defined as the training datasets. The normalized root-mean-squared error expressed as a percentage (FitPercent^47^) was employed as a measure of how well the model response fit the validation dataset. For each validation round, FitPercent was calculated for the cell density or fatty acid concentration as follows:

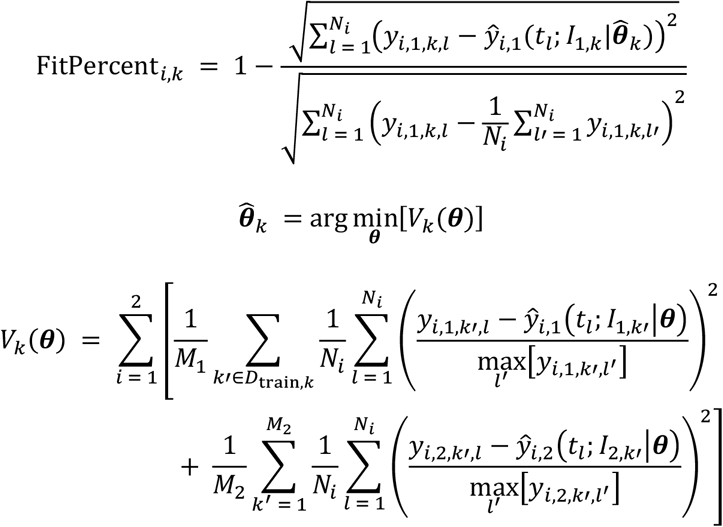

The subscript *k* represents the batch of strain FA2 selected for validation dataset. The subscript set *D*_train,*k*_ represents the set of FA2 strain batches selected for the training dataset corresponding to *k*.

### Model-based input optimization

Simulink Design Optimization was used for optimizing the IPTG input. The optimal values *I*_opt_ for the strains FA2 and FA3 were chosen to maximize the fatty acid yields obtained at the end of the culture period, as follows:

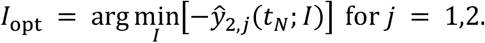

There is another decision variable for the strain FA4: the time of addition of Inducer 2 *t*_*u*_. The optimal values *I*_opt_ and *t*_*u*,opt_ for the strain FA4 were chosen as follows:

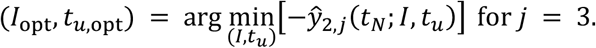

The subscript *j* represents the strain index (*j* = 1 for FA2, *j* = 2 for FA3, and *j* = 3 for FA4). The culture duration *t*_*N*_ in the simulation was defined as 25 h, as in the experiments. As with the parameter estimation, the *lsqnonlin* command was used for optimization, and the trust region method was selected as its algorithm.

## Data availability

The datasets and computer codes used in this study are available at GitHub (https://github.com/TomokiOhkubo/HISICC2).

## Acknowledgments

We thank ACS Authoring Services (https://authoringservices.acs.org/) for English language editing. This study was supported by the Next Generation Interdisciplinary Research Project of Nara Institute of Science and Technology (NAIST) and AMED under Grant Number JP23wm0425017.

## Author contributions

T.O. designed the research, performed the modeling and simulation, and wrote the manuscript. F.Z. prepared the experimental data. All authors contributed to the article and approved the final manuscript.

## Declaration of interests

The authors declare that they have no competing interests.

## Notes

### Competing Interest Statement

The authors have declared no competing interest.

