## Supporting information for "Designing a hybrid in silico/in-cell controller robust to process–model mismatch associated with dynamically regulated enzyme overexpression"

#### Author information

Tomoki Ohkubo[1*], Yuichi Sakumura[1,2], Fuzhong Zhang[3], Katsuyuki Kunida[1,4]
[1] Graduate School of Science and Technology, Nara Institute of Science and Technology, Ikoma, Nara, 8916-5, Japan
[2] Data Science Center, Nara Institute of Science and Technology, Ikoma, Nara, 8916-5, Japan
[3] Department of Energy, Environmental and Chemical Engineering, Washington University in St. Louis, St. Louis, MO 63130, USA
[4] School of Medicine, Fujita Health University, Toyoake, Aichi, 470-1192, Japan

#### Supporting information

##### Table S1.

Process outputs and measurement time in the experimental dataset

| Process output | | Measurement time | |
|---|---|---|---|
| **Index $i$** | **Name** | **Index $l$** | **Value [h]** |
| 1 | Relative cell density | 1 | 0 |
| | | 2 | 1 |
| | | 3 | 5 |
| | | 4 | 9 |
| | | 5 | 13 |
| | | 6 | 17 |
| | | 7 | 21 |
| | | 8 $(= N_1)$ | 25 |
| 2 | Fatty acid concentration | 1 | 0 |
| | | 2 $(= N_2)$ | 25 |

### Table S2.

*Escherichia coli* strains and IPTG input levels in the experimental dataset

| **Strain** | | **IPTG input level** | |
|---|---|---|---|
| **Index** $j$ | **Name** | **Index** $k$ | **Value [µM]** |
| 1 | FA2 | 1 | 0 |
| | | 2 | 10 |
| | | 3 | 40 |
| | | 4 | 100 |
| | | 5 | 400 |
| | | 6 $(= M_1)$ | 1000 |
| 2 | FA3 | 1 $(= M_2)$ | 100 |
